# An elastic-net logistic regression approach to generate classifiers and gene signatures for types of immune cells and T helper cell subsets

**DOI:** 10.1101/623082

**Authors:** Arezo Torang, Paraag Gupta, David J. Klinke

## Abstract

**Background:** Host immune response is coordinated by a variety of different specialized cell types that vary in time and location. While host immune response can be studied using conventional low-dimensional approaches, advances in transcriptomics analysis may provide a less biased view. Yet, leveraging transcriptomics data to identify immune cell subtypes presents challenges for extracting informative gene signatures hidden within a high dimensional transcriptomics space characterized by low sample numbers with noisy and missing values. To address these challenges, we explore using machine learning methods to select gene subsets and estimate gene coefficients simultaneously.

**Results:** Elastic-net logistic regression, a type of machine learning, was used to construct separate classifiers for ten different types of immune cell and for five T helper cell subsets. The resulting classifiers were then used to develop gene signatures that best discriminate among immune cell types and T helper cell subsets using RNA-seq datasets. We validated the approach using single-cell RNA-seq (scRNA-seq) datasets, which gave consistent results. In addition, we classified cell types that were previously unannotated. Finally, we benchmarked the proposed gene signatures against other existing gene signatures.

**Conclusions:** Developed classifiers can be used as priors in predicting the extent and functional orientation of the host immune response in diseases, such as cancer, where transcriptomic profiling of bulk tissue samples and single cells are routinely employed. Information that can provide insight into the mechanistic basis of disease and therapeutic response. The source code and documentation are available through GitHub: https://github.com/KlinkeLab/ImmClass2019.

## Background

Host immune response is a coordinated complex system, consisting of different specialized cell types of innate and adaptive immune cells that vary dynamically and in different anatomical locations. As shown in Fig. 1, innate immune cells comprise myeloid cells including eosinophils, neutrophils, basophils, monocytes, and mast cells. Adaptive immune cells are mainly B lymphocytes and T lymphocytes that specifically recognize different antigens [1]. Linking innate with adaptive immunity are antigen presenting cells, like macrophages and dendritic cells, and Natural Killer cells. Traditionally, unique cell markers have been used to characterize and separate different immune cell subsets from heterogeneous cell mixtures using flow cytometry [2, 3, 4]. However, flow cytometry measures on the order of 10 parameters simultaneously and relies on prior knowledge for selecting relevant molecular markers, which could provide a biased view of the immune state within a sample [5]. Recent advances in technology, like mass cytometry or multispectral imaging, have expanded the number of molecular markers, but the number of markers used for discriminating among cell types within a sample remains on the order of 10^1.5^

**Figure 1.**
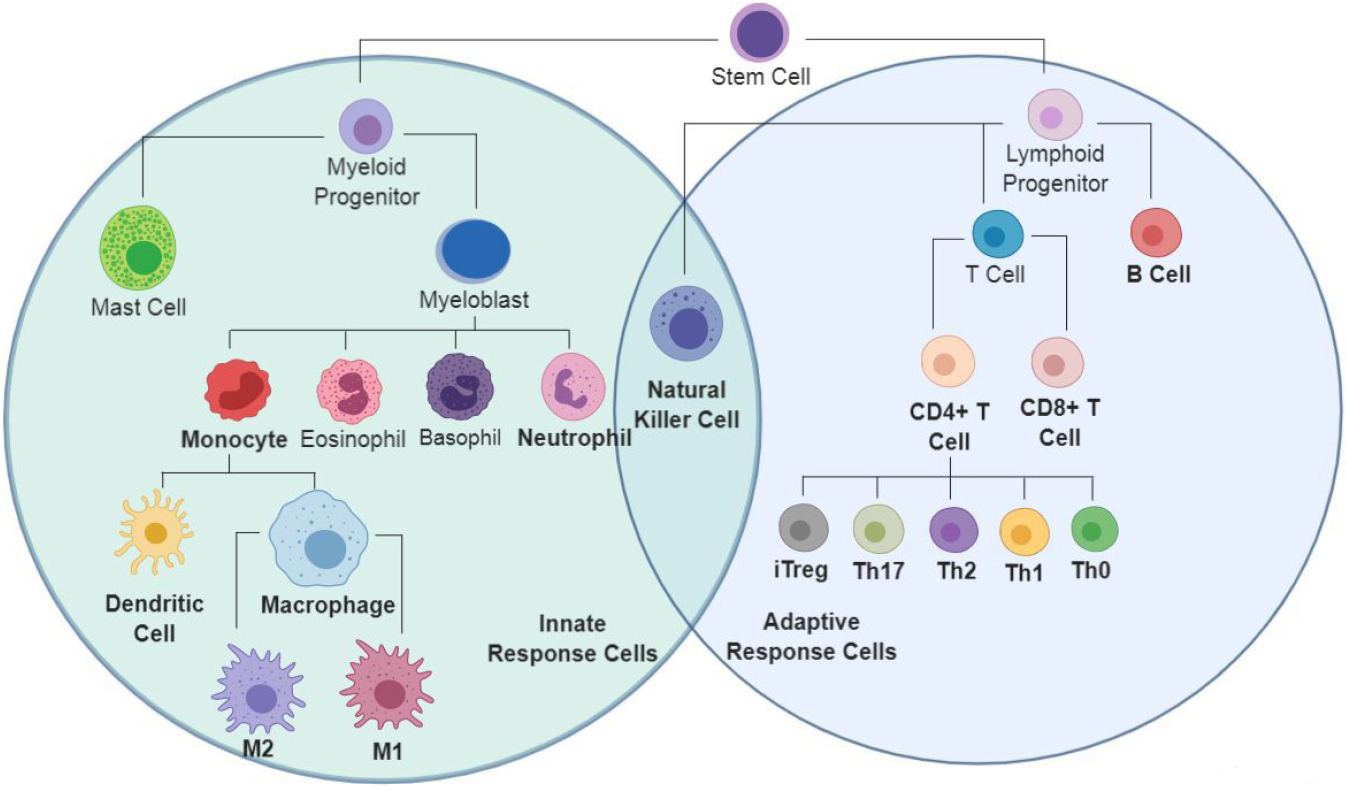
Lineage tree representation of cells of the immune system. Gene signatures were developed in this study for immune cells highlighted in bold.

In the recent years, quantifying tumor immune contexture using bulk transcriptomics or single-cell RNA sequencing data (scRNA-seq) has piqued the interest of the scientific community [6, 7, 8, 9, 10]. Advances in transcriptomics technology, like RNA sequencing, provide a much higher dimensional view of which genes are expressed in different immune cells (i.e., on the order of 10^3^) instead of focusing on a small number of genes [11]. Conceptually, inferring cell types from data using an expanded number of biologically relevant genes becomes more tolerant to non-specific noise and non-biological differences among samples and platforms. In practice, cell types can be identified using gene signatures, which are defined as sets of genes linked to common downstream functions or inductive networks that are co-regulated [12, 13], using approaches such as Gene Set Enrichment Analysis (GSEA) [12]. However, as microarray data can inflate detecting low abundance and noisy transcripts and scRNA-seq data can have a lower depth of sequencing, opportunities for refining methods to quantify the immune contexture using gene signatures still remain.

Leveraging transcriptomics data to identify types of immune cells presents analytic challenges for extracting informative gene signatures hidden within a high dimensional transcriptomics space that is characterized by low sample numbers with noisy and missing values. Typically, the number of cell samples is in the range of hundreds or less, while the number of profiled genes is in the tens thousands [14]. Yet, only a few number of genes are relevant for discriminating among immune cell subsets. Datasets with a large number of noisy and irrelevant genes decrease the accuracy and computing efficiency of machine learning algorithms, especially when the number of samples are very limited. Hence, it is essential to use feature selection algorithms to reduce redundant genes [15]. The application of feature selection methods enables developing gene signatures in different biomedical fields of study [16]. There are many proposed feature selection methods to select gene sets with the properties that enable high accuracy classification. In recent years, regularization methods have became more popular, which efficiently select features [17] and also control for overfitting [18]. As a machine learning tool, logistic regression is considered to be a powerful discriminative method [18]. However, logistic regression alone is not applicable for high-dimensional cell classification problems [19]. Regularized logistic regression, in the other hand, has been shown to be successfully applicable for high-dimensional problems [20]. Regularized logistic regression selects a small set of genes with strongest effects on the cost function [17]. A regularized logistic regression can be applied with different regularization terms. The most popular regularized terms are LASSO, Ridge [21], and elastic-net [22] which impose the *l*1 norm, *l*2 norm, and linear combination of *l*1 norm and *l*2 norm regularization, respectively, to the cost function. It has been shown that, specially in very high dimensional problems, elastic-net outperforms LASSO and Ridge [17, 22].

In this study, we focused on two-step regularized logistic regression techniques to develop immune cell signatures and immune cell and T helper cell classifiers using RNA-seq data for the cells highlighted in bold in Fig. 1. The first step of the process included a pre-filtering phase to select the optimal number of genes and implemented an elastic-net model as a regularization method for gene selection in generating the classifiers. The pre-filtering step reduced computational cost and increased final accuracy by selecting the most discriminative and relevant set of genes. Finally, we illustrate the value of the approach in annotating gene expression profiles obtained from single-cell RNA sequencing. The second step generated gene signatures for individual cell types using selected genes from first step and implemented a binary regularized logistic regression for each cell type against all other samples.

## Results

We developed classifiers for subsets of immune cells and T helper cells separately with two main goals. First, we aimed to annotate RNA-seq data obtained from an enriched cell population with information as to the immune cell identity. Second, we developed gene signatures for different immune cells that could be used to quantify the prevalence from RNA-seq data obtained from a heterogeneous cell population. Prior to developing the classifiers, the data was pre-processed to remove genes that have low level of expression for most of samples (details can be found in Methods section) and normalized to increase the homogeneity in samples from different studies and to decrease dependency of expression estimates to transcript length and GC-content. Genes retained that had missing values for some of the samples were assigned a values of −1. Next, regularized logistic regression (elastic-net) was performed and the optimal number of genes and their coefficients were determined.

### Generating and validating an immune cell classifier

In development of the immune cell classifier, we determined the optimal number of genes in the classifier by varying the lambda value used in the regularized logistic regression of the training samples and assessing performance. To quantify the performance using different lambdas, a dataset was generated by combining true-negative samples, which were created by randomly scrambling associated genes and their corresponding value from the testing datasets, with the original testing data, which were untouched during training and provided true-positive samples. The accuracy of predicting the true-Positive samples were used to generate Receiver Operating Characteristic (ROC) curves (Fig. 2a). Performance using each lambda was quantified as the Area Under the ROC Curve (AUC).

**Figure 2.**
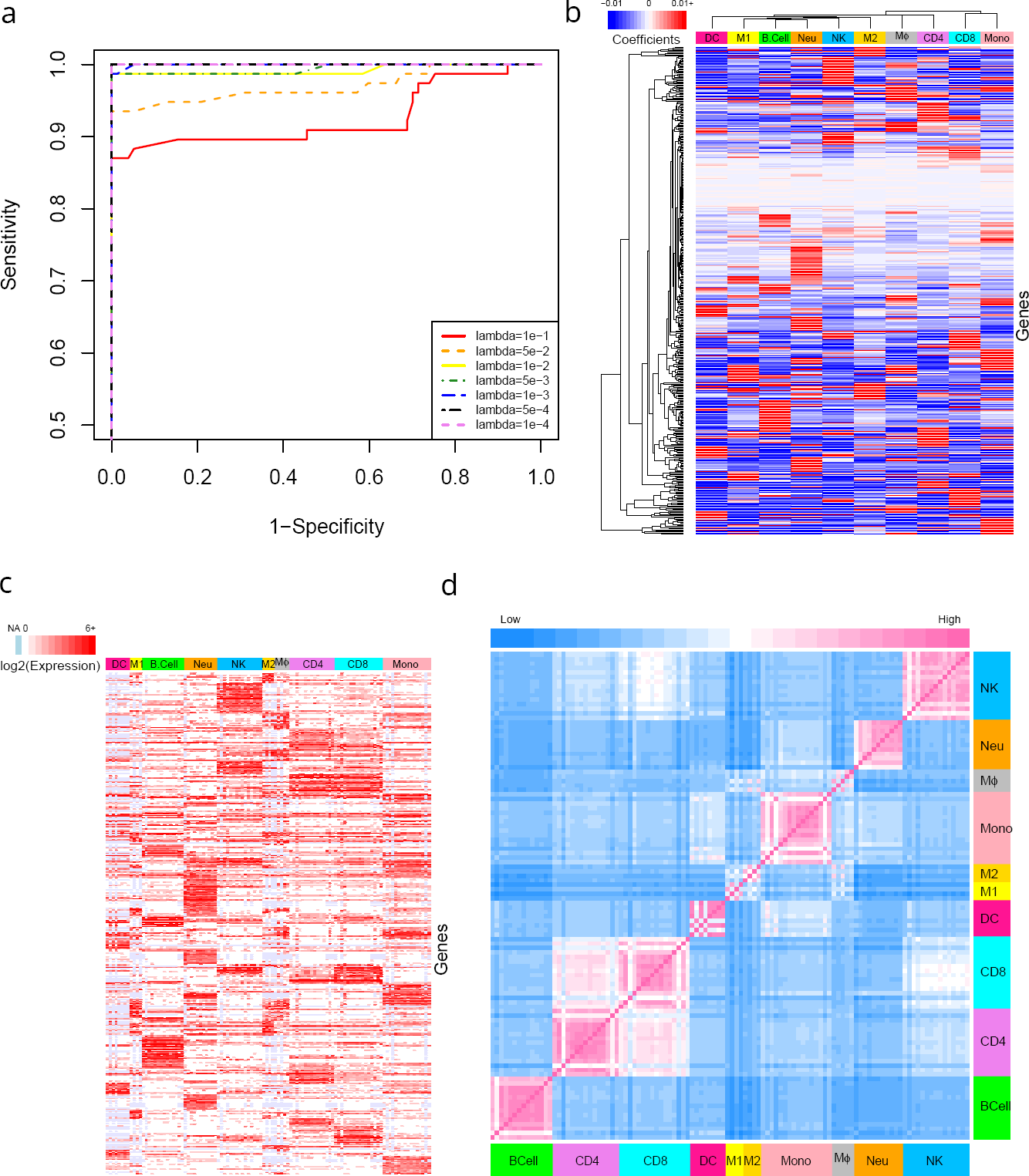
Development of immune cell classifier and similarity heatmap. a) ROC curve for the immune cell classifier was calculated using the indicated lambda values and 10-fold cross validation. The lambda value that maximized the AUC value was used for subsequent calculations. Elastic-net logistic regression was used 1to discriminate among ten immune cell types, where the value of the non-zero coefficients (panel b), expression levels (panel c), and similarity map (panel d) for the 452 genes included in the classifier are indicated by color bars. In panel b, blue to red color scheme indicates coefficients ranging from negative to positive values. Ordering of genes is the same in panels b and c. Similarity between samples calculated using distance matrix based on same 452 genes.

The optimal lambda for immune cell classifier was the smallest value (i.e., highest number of genes) that maximized the AUC. Functionally, this lambda value represents the trade-off between retaining the most possible number of informative genes (i.e., classifier signal) in the first step for developing the gene signature later, while not adding non-informative genes (i.e., classifier noise). Consequently, we selected a lambda value of 1e-4 (452 genes) for the immune cell classifier, where the selected genes and their coefficients are shown in Table S1.

To explore correlations between the weights of selected genes with their expression level, we generated heatmaps shown in Fig. 2, panels b and c. A high level of gene expression is reflected as a larger positive coefficient in a classifier model, while low or absent expression results in a negative coefficient. This is interpreted as, for example, if gene A is not in cell type 1, the presence of this gene in a sample decreases the probability for that sample to be cell type 1. For instance, E-cadherin (CDH1) was not detected in almost all monocyte samples and thus has a negative coefficient. Conversely, other genes are only expressed in certain cell types, which results in a high positive coefficient. For instance, CYP27B1, INHBA, IDO1, NUPR1, and UBD are only expressed by M1 macrophages and thus have high positive coefficients.

The differential expression among cell types suggests that the set of genes included in the classifier model may also be a good starting point for developing gene signatures, which is highlighted in Fig. 2d. Here, we focused on the expression of the 452 genes included in the classifier model and the correlations between samples clustered based on cell types. The off-diagonal entries in the correlation matrix are colored by euclidean distance values with the color indicating similarity between sample pairs (similar: pink versus dissimilar: blue) and color bars along the axes highlight the cell types for the corresponding RNA-seq samples. As expected, RNA-seq samples from the same cell type were highly similar. More interestingly, correlation between different cell types can also be seen, like high similarity between CD4+ and CD8+ T cell samples, CD8+ T cell and NK cell samples, and monocyte and dendritic cell samples. Collectively, these heatmaps illustrate that the selected genes are a highly condensed but still representative set of genes that include main characteristics of the immune cell types. It is also notable to compare the clustering result of cell types based on their coefficients in the classifier shown in Fig. 2b with similarity matrix in Fig. 2d. Since in the classifier coefficients are forcing the model to separate biologically close cell types (like CD4+ T cell and CD8+ T cell), the resulted clustering did not find them in close relationship (Fig. 2b). However, in the case of their expression values, their similarity is remains (Fig. 2d).

### Evaluating the Immune Cell classifier using scRNA-seq datasets

To evaluate the proposed classifier in immune cell classification, two publicly accessible datasets generated by scRNA-seq technology were used [23, 24]. The first dataset reported by [23] included malignant, immune, stromal and endothelial cells from 15 melanoma tissue samples. We focused on the immune cell samples, which included 2761 annotated samples of T cells, B cells, M*phi* and NK cells, and 294 unresolved samples. The immune cells in this study were recovered by flow cytometry by gating on CD45 positive cells. Annotations were on the basis of expressed marker genes while unresolved samples were from the CD45-gate and classified as non-malignant based on inferred copy number variation (CNV) patterns (i.e., CNV score *<* 0.04).

Following a pre-processing step to filter and normalize the samples similar to the training step, the trained elastic-net logistic regression model was used to classify cells into one of the different immune subsets based on the reported scRNA-seq data with the results summarized in Fig. 3a. The inner pie chart shows the prior cell annotations reported by [23] and the outer chart shows the corresponding cell annotation predictions by our proposed classifier. Considering T cells as either CD4+ T cell or CD8+ T cell, the overall similarity between annotations provided by [23] and our classifier prediction is 96.2%. The distribution in cells types contained within the unresolved samples seemed to be slightly different than the annotated samples as we predicted the unresolved samples to be mainly CD8+ T cells and B cells.

**Figure 3.**
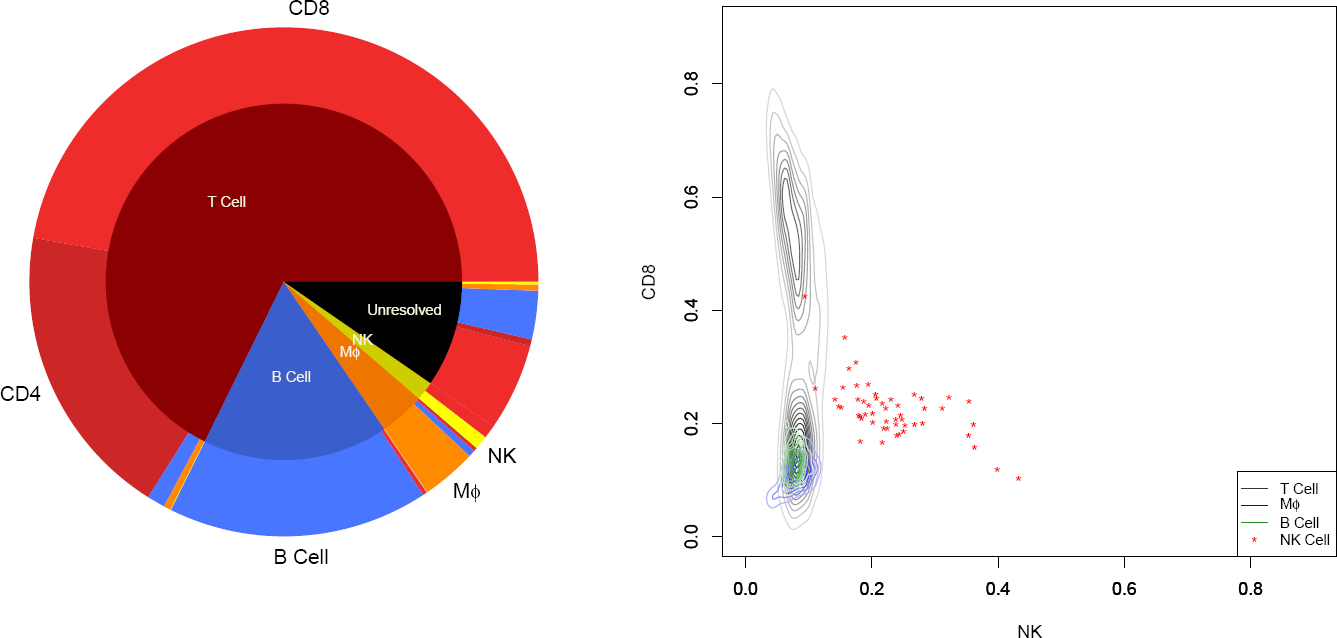
Immune cell annotation prediction based on scRNA-seq data against prior annotations reported in melanoma dataset. a) The inner pie chart summarizes the cell annotations reported by Tirosh et al [23] and includes 298 unannotated CD45-positive non-malignant cells (labeled as Unresolved) isolated from melanoma tissue samples. Unannotated samples were acquired following gating for CD45+ single cells and classified as non-malignant based on inferred copy number variation patterns. Using gene expression values reported for each scRNA-seq sample, a new cell annotation was determined based on the closest match with the alternative cell signatures determined using elastic-net logistic regression, which are summarized in outer pie chart. b) The contour plot for the likelihood of a sample to be either an NK cell or CD8+ T cell based on gene expression stratified by cells previously annotated by [23] to be T cells, macrophages, B cells, or NK cells.

The only cell type with low similarity between our classifier predictions and prior annotations was NK cells, where we classified almost half of samples annotated previously as NK cells as CD8+ T cell. Discriminating between these two cell types is challenging as they share many of the genes related to cytotoxic effector function and can also be subclassified into subsets, like CD56bright and CD56dim NK subsets [25]. To explore this discrepancy, we compared all annotated samples based on their CD8 score and NK score provided by the classifier, as shown in Fig. 3b. Although the number of NK cell samples are relatively low, it seems that the NK samples consist of two groups of samples: one with a higher likelihood of being a NK cell and a second with almost equal likelihood for being either CD8+ T cell or NK cell. We applied principal component analysis (PCA) to identify genes associated with this difference and used Enrichr for gene set enrichment [26, 27]. Using gene sets associated with the Human Gene Atlas, the queried gene set was enriched for genes associated with CD56 NK cells, CD4+ T cell and CD8+ T cell. Collectively, the results suggests that the group of cells with similar score for NK and CD8 in the classifier model are Natural Killer T cells.

We also analyzed a second dataset that included 317 epithelial breast cancer cells, 175 immune cells and 23 non-carcinoma stromal cells, from 11 patients diagnosed with breast cancer [24]. We only considered samples annotated previously as immune cells, which were annotated as T cells, B cells, and myeloid samples by clustering the gene expression signatures using non-negative factorization. The scRNA-seq samples were similarly pre-processed and analyzed using the proposed classifier, with the results shown in Fig. 4. The inner pie chart shows the prior cell annotations reported by [24] and the outer chart shows the corresponding predicted cell annotation by our proposed classifier. Considering T cells as either CD4+ T cell or CD8+ T cell, 94.4% of reported T cells are predicted as the same cell type and other 5.6% is predicted to be DC or NK cells. However, for reported B cells and myeloid cells, we predicted relatively high portion of samples to be T cells (15.7% of B cells and 40% of myeloid cells). The rest of the myeloid samples were predicted to be macrophages or dendritic cells. Collectively, our proposed classifier agreed with many of the prior cell annotations and annotated many of the samples that were previously unresolved.

**Figure 4.**
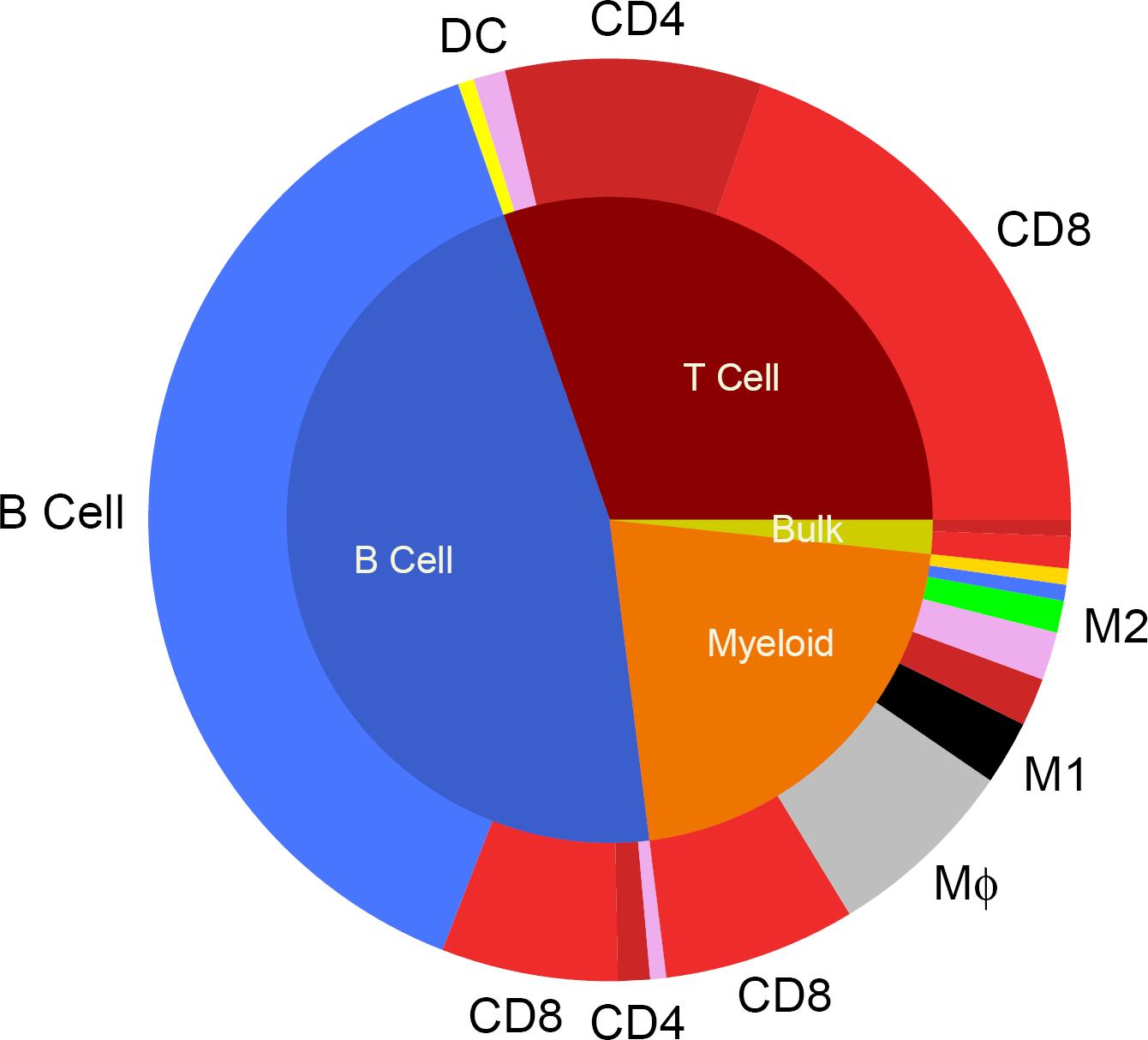
Immune cell annotation prediction against prior annotations reported in breast cancer scRNA-seq dataset. The inner pie chart summarizes the cell annotations reported by Chung et al [24], which annotated scRNA-seq results by clustering by gene ontology terms using likelihood ratio test. Using the gene expression profile reported for each scRNA-seq sample, a new cell annotation was determined based on the closest match with the alternative cell signatures determined using elastic-net logistic regression, which is summarized in the outer pie chart.

### Developing a classifier for T Helper cell subsets

Similar to the immune cell classifier, we next wanted to generate a classifier to distinguish among T helper cells and applied regularized logistic regression to corresponding training samples. We explored different values of the regression parameter lambda to find the optimal number of genes. To visualize the performance of different lambdas, we generated True-Negative samples by randomly scrambling testing datasets. Original testing data that were completely untouched during training were used as True-Positive samples. The True-Negative and True-Positive samples were used to generate ROC curves (Fig. 5a) and the AUC was used to score each lambda value. Generally, the lambda values for T helper cell classifier represents the trade-off between retaining genes and keeping the AUC high. However, there appeared to be an inflection point at a lambda value of 0.05 whereby adding additional genes, by increasing lambda, reduced the AUC. Consequently, we selected a lambda value equal to 0.05 (72 genes) for T helper classifier. The selected genes and their coefficients are listed in Table S1. The gene list was refined subsequently by developing a gene signature.

**Figure 5.**
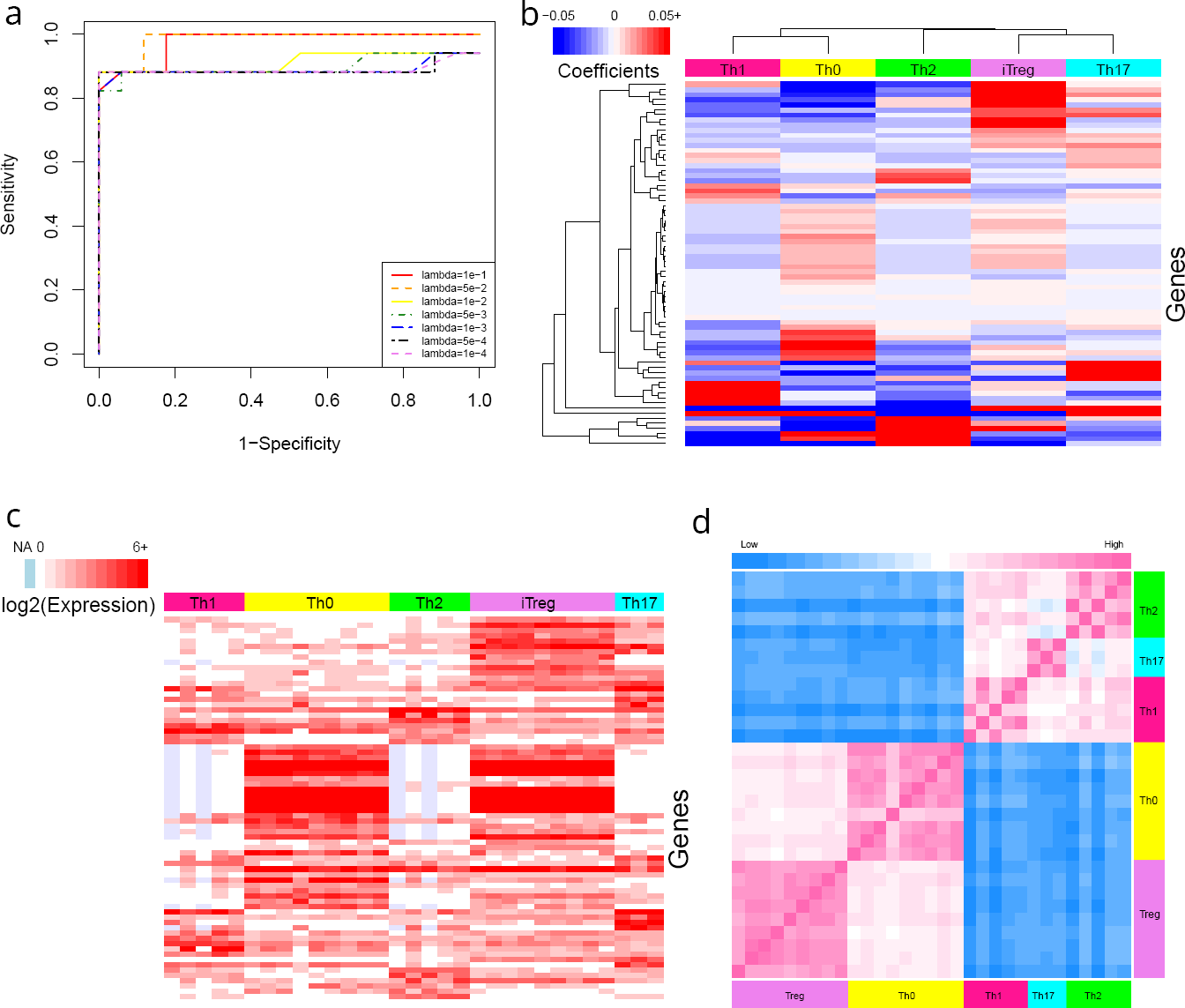
Development of T helper cell classifier and similarity heatmaps. a) ROC curve for the T helper cell classifier was calculated using the indicated lambda values and 10-fold cross validation. The lambda value that maximized the AUC value was used for subsequent calculations. Elastic-net logistic regression to discriminate among five T helper cell types, where the value of the non-zero coefficients (panel b), expression levels (panel c), and similarity map (panel d) for the 72 genes included in the classifier are indicated by color bars. In panel b, blue to red color scheme indicates coefficients ranging from negative to positive values. Ordering of the genes is the same in panels b and c. In panel d, similarity between samples calculated using a euclidean distance matrix based on same 72 genes, where the color indicates the distance (pink: high similarity/low distance; blue: low similarity/high distance). Color bar on the top/side of the heatmap indicates the cell type of origin.

Similar to the immune cell classifier, the coefficients of the selected genes for the T helper cell classifier correlated with their expression levels, as seen by comparing the heatmaps shown in Fig. 5, panels b and c. For instance, FUT7 has been expressed in almost all T helper cell samples except for iTreg that result in a negative coefficient for this cell type. In addition, there are sets of genes for each cell type that have large coefficients only for certain T helper cell subsets, like ALPK1, TBX21, IL12RB2, IFNG, RNF157 for Th1 that have low expression in other cells. As illustrated in Fig. 5d, the genes included in the classifier don’t all uniquely associate with a single subset but collectively enable discriminating among T helper cell subsets. Interestingly, the T helper subsets stratified into two subgroups where naive T helper cells (Th0) and inducible T regulatory (iTreg) cells were more similar than effector type 1 (Th1), type 2 (Th2), and type 17 (Th17) T helper cells. Similar to the immune cell classifier, we also noted that the clustering of the classifier coefficients is different from what similarity matrix shows in Fig. 5d because the classifier coefficients aim to create a “classifying distance” among closely related cell types.

Finally by comparing the results of immune cell classifier with that of the T helper classifier, the intensity of differences among cell types can be seen in Fig. 2c and Fig. 5c. In the first figure you can find completely distinct set of genes in each cell type while in the second figure the gene sets are not as distinct which could be due to either the few number of samples or high biological similarity between T helper cell types.

### Application of the Classifiers

Clinical success of immune checkpoint inhibitors (ICI) for treating cancer coupled with technological advances in assaying the transcriptional signatures in individual cells, like scRNA-seq, has invigorated interest in characterizing the immune contexture within complex tissue microenvironments, like cancer. However as illustrated by the cell annotations reported by [24], identifying immune cell types from noisy scRNA-seq signatures using less biased methods remains an unsolved problem. To address this problem, we applied our newly developed classifiers to characterize the immune contexture in melanoma and explored differences in immune contexture that associated with immune checkpoint response. Of note, some melanoma patients respond to ICIs durably but many others show resistance [28]. Specifically, we annotated immune cells in the melanoma scRNA-seq datasets [23, 29] using our classifiers separately for each patient sample and ordered samples based on the treatment response, with the results shown in Fig. 6a, b. We used the percentage of cell type for each tumor samples as it is more informative and meaningful than using absolute cell numbers. It is notable that untreated and NoInfo samples likely include both ICI-resistant and ICI-sensitive tumors.

**Figure 6.**
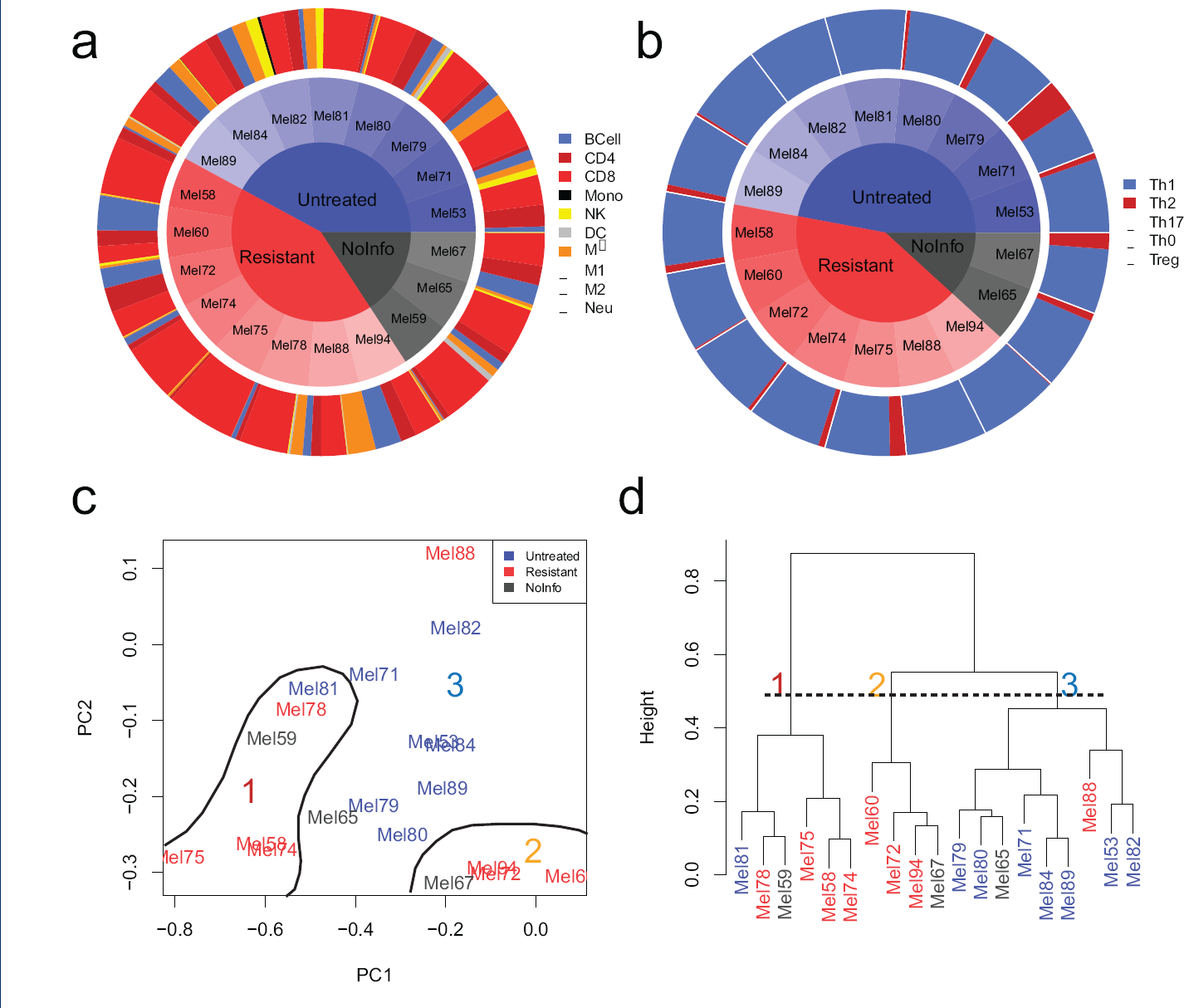
Annotation of scRNA-seq results from melanoma dataset stratified by patient treatment status. Treatment status of patients diagnosed with melanoma was stratified based on their response to ICIs ([23, 29]). a) The distribution in immune cell annotations and b) T helper cell annotations based on scRNA-seq data were separated into samples obtained from ICI-resistant tumors, untreated tumors, and tumors reported in melanoma data without information about treatment status. Cell annotations were based on immune cell classifier and T helper cell classifier results. c) PCA analysis was applied on data obtained from both classifiers. d) Samples were clustered based on the percentages of nine immune cells and five T helper cells

In comparing samples from resistant tumors to untreated tumors, we found interestingly that there are samples with high prevalence of NK in untreated tumors (Mel53, Mel81, and Mel82) while no samples in resistant tumors have a high prevalence of NK cells. The mentioned untreated tumors also have no or very low number of Th2 cells in their populations. In addition, untreated tumors have a more uniform distribution of immune cell types in contrast to ICI-resistant ones, which could reflect a therapeutic bias in immune cell prevalence in the tumor microenvironment due to ICI treatment.

Next, we combined the annotation data from both classifiers and applied PCA and clustering analysis, as shown in Fig. 6, panels c and d. Using scrambled data to determine principal components and their associated eigenvalues that are not generated by random chance (i.e., a negative control), we kept the first and second principal components that capture 68% and 21% of the total variance, respectively, and neglected other components that fell below the negative control of 8.4%. As it shown in 6c, resistant samples mainly located in lowest value of second principal component (PC2). Upon closer inspection of the cell loadings within the eigen-vectors, the low values of PC2 corresponds to a low prevalence of M*ϕ* or high percentage of B cells. In addition, based on the first principal component (PC1), resistant samples have either lowest values of PC1 (Mel74, Mel75, Mel58, Mel 78) which correspond to higher than average prevalence of CD8+ T cells or highest values of PC1 (Mel60, Mel72, Mel94) that show higher than average prevalence of B cells.

In hierarchical clustering, the optimal number of clusters was selected based on calculation of different cluster indices using the NbClust R package [30] which mainly identified two or three clusters as the optimal number. In considering three groupings of the hierarchical clustering results shown in 6d, seven out of eight ICI-resistant samples clustered in first two clusters while the third cluster mainly contained untreated samples. The comparison of results from PCA and clustering analyses shows that the first cluster contained samples with extreme low value of PC1 which itself divided into two groups; one with extreme low value of PC2 and the other with higher amount of PC2. The second cluster located in highest amount of PC1 and lowest amount of PC2. All remained samples were clustered as third group, which were predominantly untreated samples. The difference in clustering suggests dissimilarities between ICI-resistant and untreated samples and the possibility of having ICI-sensitive tumors in untreated samples.

### Developing Gene Signatures

While classifiers are helpful for annotating scRNA-seq data as the transcriptomic signature corresponds to a single cell, gene signatures are commonly used to determine the prevalence of immune cell subsets within transcriptomic profiles of bulk tissue samples using deconvolution methods. Leveraging the classifier results, we generated corresponding gene signatures using binary elastic-net logistic regression. Specifically, classifier genes with non-zero coefficients were used as initial features of the models, which were regressed to the same training and testing datasets as used for developing the classifiers. Lambda values were selected for each immune and T helper cell subset based on similar method of lambda selection for classifiers and their values and corresponding AUC are shown in Table S2. Finally, all generated signatures are summarized in Table S3.

We visualized the expression levels of remained set of genes, which at least occur in one gene signature, in Fig. 7. The expression of genes retained in immune cell signatures (Fig. 7a) and T helper cell signatures (Fig. 7b) were clustered by similarity in expression (rows) and by similarity in sample (columns). For both immune and T helper cell subsets, samples of same cell type were mainly clustered together. The only exception is for macrophages (M*ϕ* and M2) which can be attributed to high biological similarity and a low number of technical replicates for these cell types.

**Figure 7.**
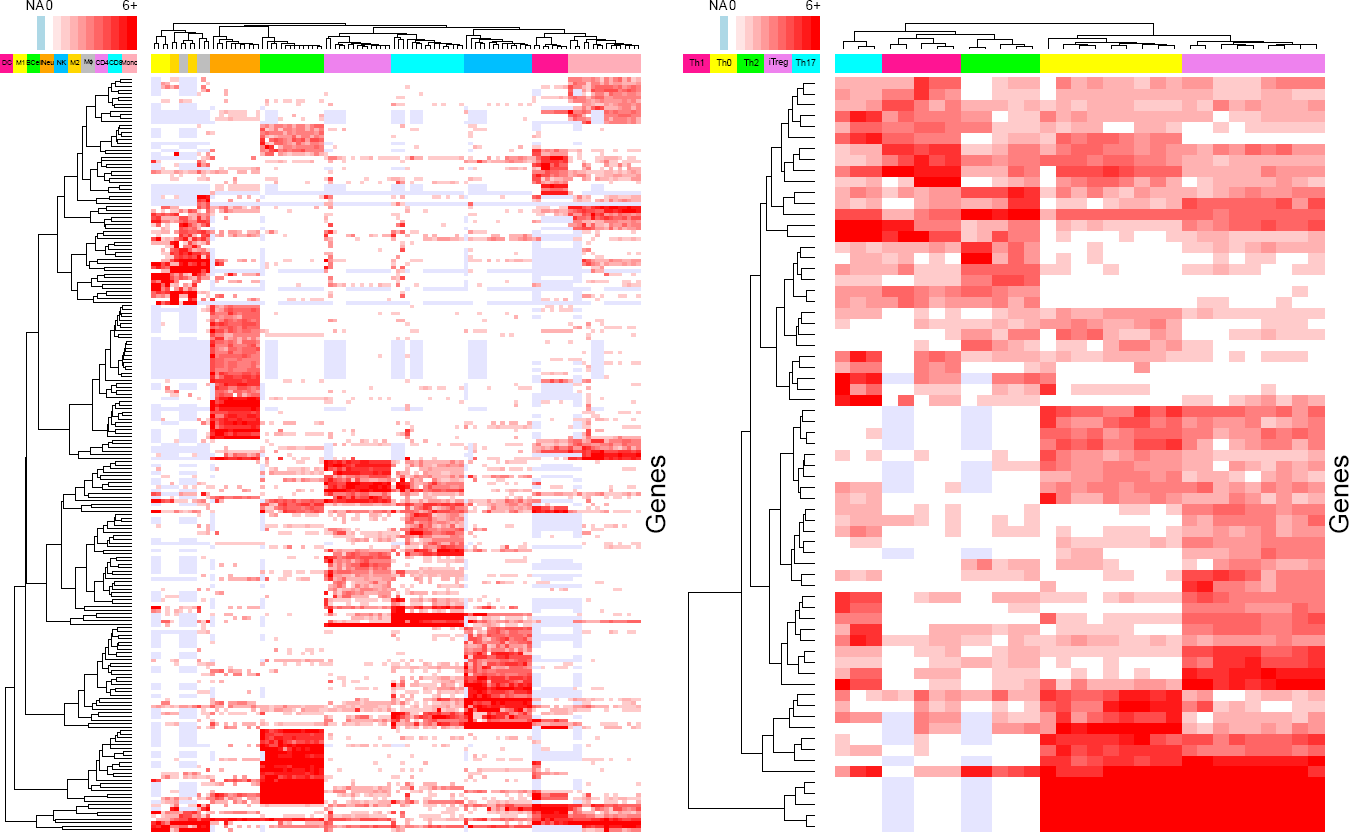
Heatmaps of the expression levels for the final list of genes created by gene signatures. The expression of genes retained in immune cell signatures (panel a) and T helper cell signatures (panel b) were clustered by similarity in expression (rows) and by similarity in sample (columns) The color bar at the top indicates the sample cell type.

In general, the gene set generated from the logistic regression model performed well with far fewer requisite genes in the testing set, a desirable result for a gene set intended to be used for immunophenotyping. In Fig. 8, the results of the bench-marking are shown separated by comparative gene set. Both the CIBERSORT and Single-Cell derived gene sets contain an average of 64 and 135 genes, respectively, while the logistic regression gene set contains an average of just 19. The new logistic regression gene set performed comparably to the existing contemporary gene sets and far exceeded the performance of the manually curated gene set used previously [6]. The benchmarking results indicate that logistic regression gene set is an improvement in efficacy over compact gene sets, such as those that are manually annotated or hand-picked. Meanwhile, the logistic regression gene set also demon-strates an optimization of broader gene sets that contain too many genes for deep specificity when used in further analysis. The inclusion of too many genes in the set can dilute the real data across a constant level of noise, while including too few lacks the power to draw conclusions with high confidence. The logistic regression gene set demonstrates a balance of these two issues through its highly refined selection of genes that can be fine-tuned using its lambda parameter.

**Figure 8.**
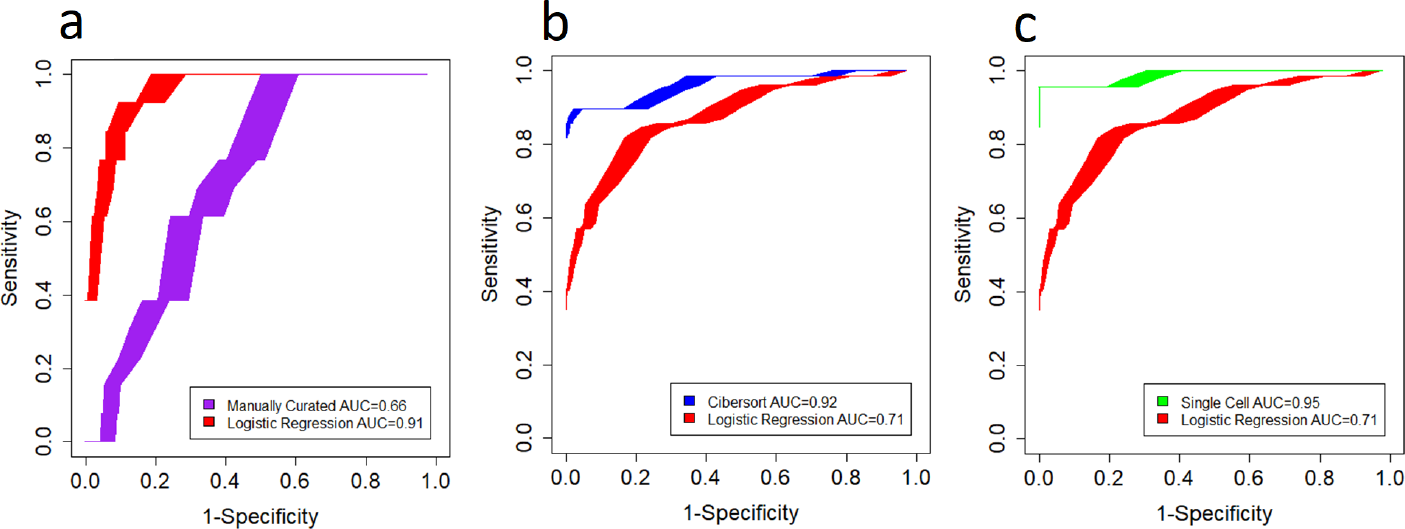
Benchmarking ROC performance curves. ROC curves to illustrate relative performance between logistic regression gene set and the manually curated (Panel A), CIBERSORT (Panel B), and single cell gene sets (Panel C). The logistic regression gene set’s performance is shown in red. Shaded regions are 95% confidence intervals about the average ROC curve simulated from 1000 iterations.

## Discussion

Recent developments in RNA sequencing enable a high fidelity view of the transcriptomic landscape associated with host immune response. Despite considerable progress in parsing this landscape using gene signatures, gaps remain in developing unbiased signatures for individual immune cell types from healthy donors using high dimensional RNA-seq data. Here, we developed two classifiers - one for immune cell subsets and one for T helper cell subsets - using elastic-net logistic regression with cross validation. The features of these classifiers have been used as starting point for generation of gene signatures captured with fifteen binary elastic-net logistic regression models as the most relevant gene sets to distinguish among different immune cell types without making too much noise.

Gene signatures in previous studies have been developed and used mainly as a base for deconvolution of tumor microenvironment and to find the fractions of existing immune cells. Therefore, as the first step, determining cell-specific gene signatures critically influences the results of deconvolution methods [31]. Newman et al. defined gene signatures for immune cells using two-sided unequal variances t-test as base matrix for CIBERSORT [8]. In another study, Li et al. in developing TIMER, generated gene signatures for six immune cell types with selecting genes with expression levels that have a negative correlation with tumor purity [9]. More recently, Racle et al. developed a deconvolution tool based on RNA-seq data (EPIC) by pre-selecting genes based on ranking by fold change and then selected genes by manually curating and comparing the expression levels in blood and tumor microenvironment [10]. Finally, quanTIseq (the most recently developed tool for deconvolution) has been developed for RNA-seq data based on the gene signatures generated by quantizing the expression levels into different bins and selecting high quantized genes for each cell type that have low or medium expression in other cell types [7]. Although all methods obtained high accuracy based on their developed signatures, a more rigorous and unbiased gene signature developed by RNA-seq data and precise feature selection methods can be used to improve the accuracy even further and validate the process for downstream analyses.

In addition, to identify cell types based on their transcriptome, clustering techniques have been used in many studies [32, 33]. However, there are high variability levels of gene expression even in samples from the same cell type. Moreover, transcriptomics data has high dimensions (tens of thousands) and this is too complicated for clustering techniques specially because only few number of genes are discriminative. To overcome these problems some studies used supervised machine learning methods like Support Vector Machine (SVM) [34, 35]. However, to the best of our knowledge, this paper is the first to apply two-step regularized logistic regression on RNA-seq transcriptomic of immune cells. This method increases the chance to capture the most discriminative set of genes for each cell type based on the power of an elastic-net [22]. In addition, using a two-step elastic net logistic regression enabled eliminating the most irrelevant genes while keeping the most possible significant genes in the first step and more deeply selecting among them in the second step to generate robust gene signatures for immune cells.

Moreover, contemporary methods have only considered a limited number of immune cell types, and specifically T helper subsets as individual cell types have been neglected [23, 29, 24] in comprehensive studies. Therefore, the other novel aspect of this study is the separation of models for immune cells and T helper cells and development of gene signatures for vast number of immune cell types (fifteen different immune cell types) including different T helper cell subsets. This can be used to study immune system in different diseases in more depth. As we used publicly available RNA-seq datasets for immune cells and T helper cells, we acknowledge that our developed classifiers and gene signatures may be still constrained by the limited number of samples specifically for T helper cells. As more data describing the transcriptome of for immune cells will become accessible, one can update the classifiers and gene signatures. Despite the limited number of samples used in the approach, the developed classifiers can even be applied to completely untouched and large datasets [23, 24] that have been generated using scRNA-Seq technology which creates noisier data.

## Conclusions

Here, we developed an immune cell classifier and classifier for T helper cell subsets along with gene signatures to distinguish among fifteen different immune cell types. Elastic-net logistic regression was used to generate classifiers with 10-fold cross-validation after normalizing and filtering two separate RNA-seq datasets that were generated using defined homogeneous cell populations. Subsequently, we generated gene signatures using a second step of binary regularized logistic regression applied to the RNA-seq data using previously selected classifier genes. As an external validation, the resulting classifiers accurately identified the type of immune cells in scRNA-seq datasets. Our classifiers and gene signatures can be considered for a different downstream applications. First, the classifiers may be used to detect the type of immune cells in under explored bulks and to verify uncertainly annotated immune cells. Second, the gene signatures could be used to study tumor micro-environments and the connections of immune systems with cancer cells, which is emerging to be an important clinical question.

## Methods

### Data Acquisition

RNA-seq datasets for 15 different immune cell types including T helper cells, were obtained from ten different studies [36, 37, 38, 39, 40, 41, 42, 43, 44, 45] which were publicly accessible as part of *Gene Expression Omnibus* [46]. The list of samples is provided as Supplementary Table S1. Cell types divided into two groups: the immune cells includes B cells, CD4+ and CD8+ T cells, monocytes (Mono), neutrophils (Neu), natural killer (NK) cells, dendritic cells (DC), macrophage (M*ϕ*), classically (M1) and alternatively (M2) activated macrophages, and the T helper cells includes Th1, Th2, Th17, Th0, and Regulatory T cells (Treg). The goal was to train the gene selection model on immune cell types, and CD4+ T cell subsets (T helper cells), separately. As if these two groups of cells are analyzed together, many of the genes that potentially could be used to discriminate among T helper cell subsets might be eliminated as they overlap with genes associated with CD4+ T cells.

In short, a total of 233 samples were downloaded and divided into two sets of 185 and 48 samples, for immune cells and T helper cells, respectively. Moreover, immune cell samples have been further divided into 108 training and 77 testing samples. Numbers for T helper samples are 31 and 17, respectively. Training and testing data include samples from all studies. For a verification dataset, scRNA-seq data derived from CD45+ cell samples obtained from breast cancer [24] and melanoma [23] were used with GEO accession numbers of GSE75688 and GSE72056, respectively.

### Data Normalization

The expression estimates provided by the individual studies were used, regardless of the underlying experimental and data processing methods (Table S1). For developing individual gene signatures and cell classification models, we did not use raw data due to sample heterogeneity such as different experimental methods and data processing techniques used by different studies as well as differences across biological sources. Rather, we applied a multistep normalization process before training models. To eliminate obvious insignificant genes from our data, for immune cell samples, genes with expression values higher than or equal to five, in at least five samples have been kept, otherwise, they were eliminated from the study. However, for T helper samples, due to fewer number of samples, four samples with values higher than or equal to five were enough to be considered in the study. After first step of filtering, the main normalization step was used to decrease dependency of expression estimates to transcript length and GC-content[47, 48]. For all four sets of samples, including training and testing samples for immune cells and for T helper cells, expression estimates were normalized separately by applying *withinLaneNor-malization* and *betweenLaneNormalization* functions from EDASeq package [49] in R programming language (R 3.5.3), to remove GC-content biases and between-lane differences in count distributions [49]. After normalization, the second step of filtration, just similar to the first step, was applied to eliminate genes with insignificant expression.

### Missing Values

In contrast to previous studies that only considered intersection genes [50], in order to avoid of deletion of discriminative genes, we tried to keep genes with high expression, as much as possible. However, for most of genes, values for some samples were not estimated. Hence, to deal with these missing values, we used an imputation method [51] and instead of mean imputation we set a dummy constant since mean imputation in this case is not meaningful and can increase error. Specifically, we generated a training set for each group of cell types, by duplicating the original training set 100 times and randomly eliminating ten percent of expression values. We next set −1 for all these missing values (both original missing values and those we eliminated) as a dummy constant because all values are positive and it is easy to be learned by the system as noise. This approach makes the system learn to neglect specific value (−1) and treat it like noise, instead of learning it as a feature of the samples.

### Classifier Training and Testing

Considering the few number of training samples in comparison with the high dimensions (15453 genes in immune cell samples and 9146 genes in the T helper samples) and to avoid both over fitting the model and adding noise to the prediction model, we used regularization with logistic regression to decrease the total number of genes and select the most discriminative set of genes. To perform gene selection, we trained a lasso-ridge logistic regression (elastic-net) model, which automatically sets the coefficients of a large number of genes to zero and pruned the number of genes as features of the classifier. We cross-validated the model by implementing cv.glmnet function with nfold=10 from glmnet package [21] in R programming language, using training sets for both groups of cell types. We normalized the gene expression values using a log2 transform over training sets to decrease the range of values that can affect the performance of the model (log2(counts+1)). In order to find the optimal number of genes, we tried 7 different lambdas and tested the results over the testing samples (*cv.glmnet(family=“multinomial”, alpha=0.93, thresh=1e-07, lambda=c(0.1, 0.05, 0.01, 0.005, 0.001, 0.0005, 0.0001), type.multinomial=“grouped”, nfolds=10)*). To select the optimal value for lambda, True-Negative samples were generated by randomly scrambling testing datasets, then we generated ROC curves and considered original testing datasets as True-Positive samples.

### Developing Gene Signatures

Genes selected by the classifier models were used as initial point to build gene signatures. In this case, we trained a new binary elastic-net model for each cell type by considering a certain cell type as one class and all other cell types as another class. The training and testing samples used to build gene signatures were the training and testing samples used in developing the classifiers with the difference being that they only contained the selected genes. Similar steps including dealing with missing values, applying log2 and visualization by ROC to select optimal number of genes were applied for each cell type. This two-step gene selection approach has the advantage that it eliminates a large number of undiscriminating genes at the first and finally select few number of genes for each cell type.

### Benchmarking

Fisher exact testing was used for each gene set to characterize true and systematically scrambled data as a measure of performance of the gene set as a means of distinguishing between cell subtypes. Data was scrambled by randomly redistributing expression values by gene as well as patient in order to establish negative control values for determining specificity. The threshold for expression binarization for Fisher exact testing was selected based on gene expression histograms of the data to separate the measured expression from background noise levels, with 2.48 being used as the threshold (after log2 normalization). One-thousand iterations were processed and compiled in order to produce ROC curves with 95% confidence intervals shaded about the averaged ROC curve for each gene set’s performance. The tested gene sets were the logistic regression gene set, the CIBERSORT gene set [8], the single cell gene set [29], and the manually curated gene set that had been used previously.

## List of abbreviations

ROC: receiver-operator curves
scRNA-seq: single-cell RNA-seq
AUC: area under the ROC curve
CNV: copy number variation
PCA: principal component analysis
ICI: immune checkpoint inhibitor
SVM: support vector machine

## Declarations

### Ethics approval and consent to participate

The results described in this manuscript consist of secondary analyses of existing data and was determined by the West Virginia University IRB to qualify for an exemption from human subject research under U.S. HHS regulations 45 CFR 46.101(b)(4).

### Consent for publication

All of the authors have read the final manuscript and consent for publication.

### Availability of data and material

The datasets supporting the conclusions of this article are available in Gene Expression Omnibus repository [https://www.ncbi.nlm.nih.gov] with the following GEO accession numbers: GSE60424, GSE64655, GSE36952, GSE84697, GSE74246, GSE70106, GSE55536, GSE71645, GSE66261, GSE96538, GSE75688, GSE72056. R scripts used in the analyses can be found on GitHub [https://github.com/KlinkeLab/ImmClass2019].

## Authors’ contributions

Designed study: AT and DK; performed analyses and interpreted results: AT, PG, and DK; and drafted initial manuscript: AT, PG, and DK. All authors edited and approved the final version of the manuscript.

## Additional Files

Table S1. — Coefficients of immune cell classifier and T helper cell classifier

Coefficients of immune cell classifier were located in the first sheet and coefficients of T helper cells were located in the second sheet.

Table S2. — Lambda Selection by AUC Values

Lambdas with corresponding calculated AUC. The final column shows the selected lambdas

Table S3. — Genes in developed gene signature for immune and T helper cells

Yellow boxes show genes with negative impact in possibility of being related cell type.

Table S4. — Data information used in training models.

The second sheet shows names that used in creating datasets.

